# Shifu: an integrated framework for deep learning of RNA secondary structure

**DOI:** 10.64898/2026.07.15.738762

**Authors:** Gabriel Galvez, Quentin Vicens

## Abstract

Deep learning has advanced RNA secondary-structure prediction by bypassing explicit energy rules to capture long-range dependencies, yet progress is limited less by model scale than by how structures are measured: single scores hide where and why models fail, and benchmark scores can reflect memorization of one dataset rather than genuine generalization. We address this with Shifu, a framework of three coupled parts. Shifu-Corpus is a leakage-audited dataset of 254123 sequences from six databases, with family-aware splits certified free of exact and near-duplicate leaks. The Shifu Trifecta scores a model on three axes (correctness, breadth across diverse RNAs, and whether its confidence can be trusted) rather than one number. Shifu-LMR, a family of compact RNA language models, serves as controlled experiments: changing the training corpus shifts accuracy by 0.13, and a 65-million-parameter model, Shifu-LMR-Nano, leads on correctness while running on a laptop. We release the dataset, code, and model backbones.

## Main

For over fifty years, computational methods such as dynamic programming and covariation analysis ^1,2^ have reliably predicted RNA secondary structure (the pattern of Watson–Crick base pairs) and have even helped reveal the third domain of life ^3^. Grounded in thermodynamics ^4^ and evolutionary conservation, these foundational tools remain powerful for functional inference, yet they are increasingly complemented by data-driven models that aim to overcome their longstanding thermodynamic and structural limitations.

Machine-learning approaches emerged because prediction continued to be challenged by non-Watson–Crick pairs, multi-helical junctions, pseudoknots, and multiple functional conformations ^5^. By training on known structures, these models learned to weigh structural features more accurately than energy rules alone ^6^, and deep-learning architectures, convolutional networks and transformers, pushed the field further, capturing the non-linear, long-range dependencies in a sequence and treating structure prediction as data-driven pattern recognition that bypasses explicit energy calculation ^7^.

Progress in this data-driven era is judged almost entirely by how well a model’s predicted base pairs match the true ones, summarized by the F1 score, the harmonic mean of precision and recall ^8^. The widely used micro-F1 pools every base pair into a single number, which can quietly mask weak performance on the rarer, more complex structures that a family-weighted average would expose ^8^; and both scores are all-or-nothing, counting a helix shifted by a single nucleotide, often still functional, as a complete miss. More fundamentally, any such number is only as trustworthy as the data behind it: evaluation here is not objective but relative, depending on the dataset and the split one happens to choose, so a high benchmark score is hard to read as genuine generalization rather than memorization of a particular collection.

This problem is especially acute for RNA. Unlike proteins, RNA has no large, uniformly curated structural dataset and no shared benchmark; its public data are scattered across collections built to different standards (bpRNA ^9^, ArchiveII ^10^, RNAStrAlign ^11^, RNASSTR ^12^, DSSR-derived structures ^13^ and RNAsolo ^14^) that differ by three orders of magnitude in size and in how carefully their structures are annotated. The cost is visible the moment one model is run across several of them: its accuracy can swing from very high on one collection to near-chance on another, an instability that reflects the data and the measurement rather than the model.

We call these intertwined gaps the *data problem* (corpora that are unaudited, overlapping, and skewed) and the *representation problem*: a bare string of letters does not carry the biological constraints and information that govern how RNA folds, which a model would otherwise have to rediscover, so it overfits the benchmark instead ^15,16^. RNA has not had its *AlphaFold moment* ^17^, and we argue this owes as much to data and representation as to model design. Recent work has begun to address the data side directly: RNASSTR showed that a larger, more carefully curated training corpus improves generalization to unseen RNA families ^12^, helping establish better data and shared standards as a way forward.

We address this with three coupled tools and use them to ask whether the limiting factor is the data and how it is represented rather than model scale alone, isolating each through controlled, same-architecture comparisons. Shifu-Corpus is a unified, leakage-audited dataset of 254123 RNA sequences with family-aware splits, giving the field shared, leakage-audited ground on which to train and test. The Shifu Trifecta scores a model on three complementary axes (correctness, breadth across hard, out-of-distribution data, and whether its confidence is operationally useful) rather than on a single number that can hide where a model fails. And a family of compact RNA language models (LMR, for Language Model for RNA) serves as controlled experiments that hold architecture fixed and vary one factor at a time. Together these turn an unverifiable leaderboard into an honest measurement: the same architecture swings by 0.13 in accuracy when only its training data change, and a 65-million-parameter model trained this way leads on correctness while remaining small enough to run on a laptop. What Shifu offers over prior practice is therefore not another model but a way to tell genuine progress from benchmark artifacts, and evidence that better data and representation, not parameters alone, are what will carry RNA toward its AlphaFold moment.

## Results

### A single benchmark misleads

We first asked how much a model’s measured accuracy depends on which benchmark it is measured on. Running six published external models (ERNIE-RNA ^18^, PriFold ^19^, RiNALMo ^20^, MXfold2 ^21^, UFold ^22^ and BPfold ^23^) across seven corpora (five established public sets, TR0, TS0, VL0 ^9^, ArchiveII ^10^ and bpnew ^9^, together with the test and validation splits of the leakage-audited dataset we build in the next section) reveals striking differences (Fig. 1). The same model can swing by more than 0.6 in micro-F1 between corpora, and the model that ranks first changes with the corpus. PriFold leads on four of seven sets, including TR0 at 0.98, yet drops to 0.33 on bpnew, the set built from held-out families, where BPfold instead leads at 0.77. The pattern is telling: the corpora on which the pretrained language-model baselines excel are bpRNA-derived, the same distribution they were pretrained on, so a high score there partly measures overlap with pretraining rather than generalization ^15,16^. TR0 makes this concrete: three pretrained models cluster above 0.90 on what is essentially their own training fold, while a thermodynamic hybrid and a convolutional model that were not pretrained on it score near 0.5.

**Figure 1.**
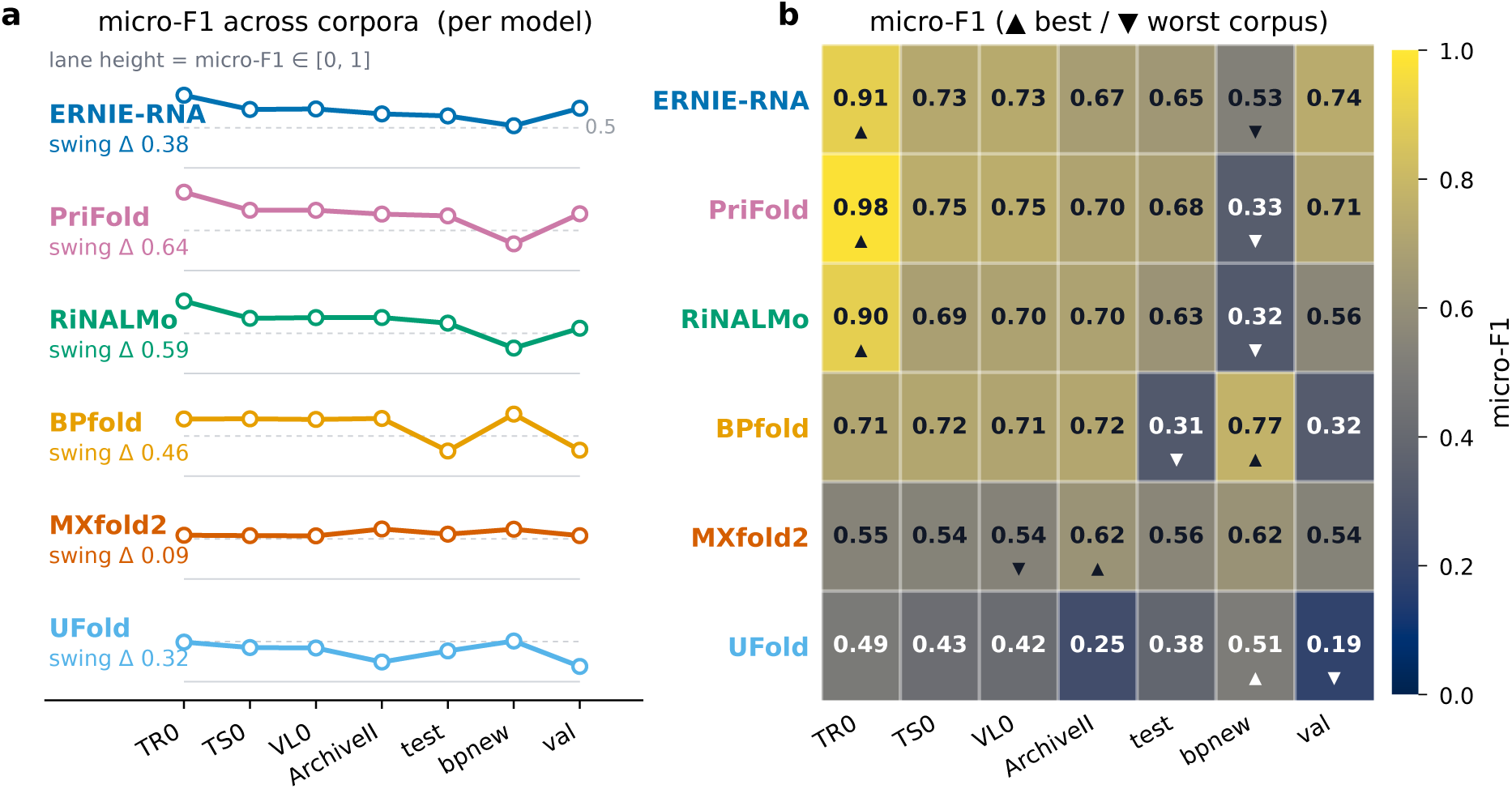
Accuracy on a single benchmark is unstable. **a**, micro-F1 of each of six external models across seven corpora (ordered by descending mean), each annotated with its swing Δ. The swing is the difference between a model’s best and worst corpus (its maximum minus its minimum micro-F1); a large Δ means a model’s measured accuracy depends heavily on which benchmark it happens to be scored on. Each lane spans micro- F1 *∈* [0, 1], and the faint dashed rule within each lane marks micro-F1 = 0.5; flat trajectories would indicate cross-corpus consistency, whereas the steep right-tail drops do not. **b**, the same model × corpus values as a heatmap, marking each model’s best (▴) and worst (V) corpus. Panels **a** and **b** are complementary views of one matrix: **a** traces each model’s trajectory across corpora (the swing), while **b**reads off the absolute values and the per-model extremes. The seven corpora are the bpRNA-1m benchmark splits TR0, TS0 and VL0^9^, the experimentally curated set ArchiveII ^10^, the held-out-family set bpnew (Rfam families absent from bpRNA training) ^9^, and the held-out test and validation splits of Shifu-Corpus (this work; Fig. 2). Sample sizes: TR0 10814, TS0 1305, VL0 1300, ArchiveII 3975, bpnew 5401, Shifu-Corpus test 25413, val 25412.

The instability is also within a benchmark, not only across benchmarks. Partitioning a single benchmark into (source × length) units (here the held-out test set of the leakage-audited dataset we build next) shows that every external model collapses to near-zero accuracy on at least one unit, typically the longest sequences, which several models cannot process at all (Extended Data Fig. 1). Two models with the same aggregate micro-F1 can therefore fail in entirely different places. The lesson from both views is the same: a single number on a single benchmark cannot tell you whether a model has learned RNA structure or just the benchmark. Two problems lie underneath, and we address them in turn: the data themselves are unaudited and overlapping, which we fix first with a leakage-audited dataset, and a single score hides where a model fails, which we then fix with a three-axis evaluation.

### Generating the Shifu dataset

Shifu-Corpus unifies six public RNA secondary-structure collections into one leakage-audited, family-split dataset (Fig. 2): no test sequence has a near-duplicate in training, and whole RNA families are held out, so measured accuracy reflects generalization rather than memorization. The six sources span the range of how RNA structures are annotated, from large bulk corpora to small hand-curated, family-stratified sets: RNASSTR^12^, bpRNA ^9^, RNAS-trAlign ^11^, ArchiveII ^10^, DSSR-derived structures ^13^ and RNAsolo ^14^. From 310034 raw input sequences, exact SHA-256 deduplication and a 20–8192 nt length filter leave 254123 unique sequences; structures are normalized to canonical Watson–Crick and wobble pairs, partitioned into source × length units, and split 80/10/10 by a family-aware procedure that keeps whole Rfam clans on one side of the split (Methods, “Family-aware splitting and leakage audit”). A leakage audit then certifies zero exact and zero near-duplicate overlap across all three pairs of splits.

**Figure 2.**
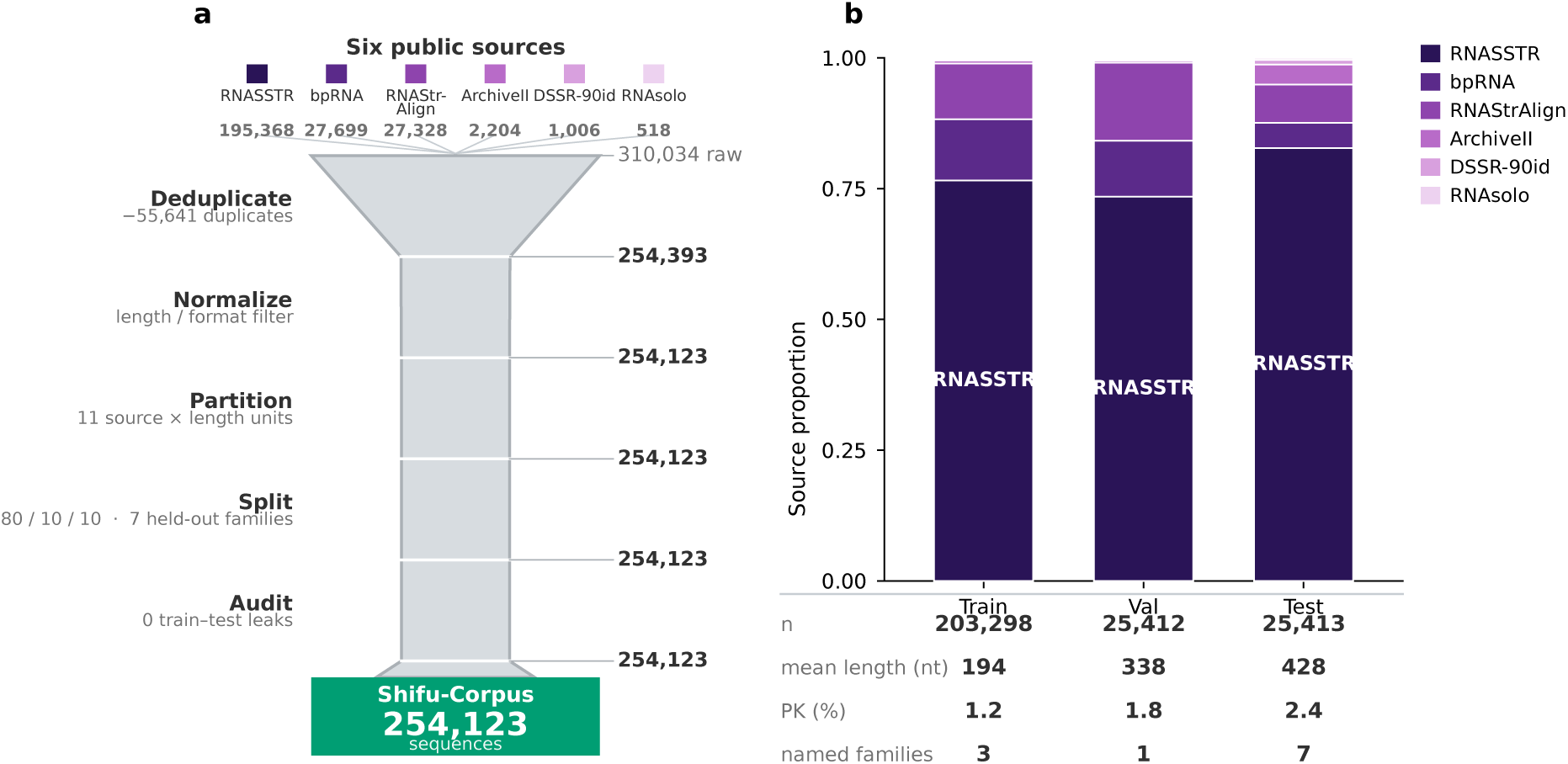
Generating the Shifu-Corpus. **a**, Construction pipeline: six public databases (with their sequence counts) are unified by exact deduplication, DSSR canonical-pair normalization, partition into source *×* length units, family-aware 80/10/10 splitting, and a leakage audit, yielding 254123 sequences (train 203298 / val 25412 / test 25413) with zero certified leaks and seven test-exclusive Rfam families. **b**, Source composition across the three splits with per-split statistics: sequence count *n*, mean length (nt), pseudoknot fraction (PK%) and the number of Rfam families. The test split is systematically longer, more pseudoknotted, and better family-labeled, so its difficulty emerges from the data rather than from subsampling.

Because sequences are grouped by family before splitting, the test set is not a re-sampling of the training data but a genuinely harder slice: its sequences are longer (mean 428 vs. 194 nt), more pseudoknotted (2.4% vs. 1.2%), and include seven Rfam families absent from training altogether (Fig. 2b). Most Shifu-Corpus sequences are unlabeled at the Rfam level, so these per-split counts reflect labeled diversity, not the full family space, which is far larger. RNASSTR contributes roughly three-quarters of the data (a property of public source sizes, not a design choice), which a source-mix penalty in the splitter keeps from saturating any one split. The result is one carefully audited dataset on which a benchmark score reflects generalization rather than memorization, offering the field a common foundation to build on. With the data on audited, leak-free ground, we can introduce the models built to test it.

### The Shifu-LMR family

The models we test on this dataset are themselves the experiment. The Shifu-LMR models (Language Model for RNA) are a controlled set of compact RNA language models that share one architecture class, training recipe and evaluation, and differ by a single design axis each, so any change in score can be assigned to that axis rather than to scale (Methods). Shifu-LMR-v0 is the full-capacity reference (291 M parameters, trained on Shifu-Corpus); against it, Shifu-LMR-Nano isolates capacity (65 M), Shifu-LMR-G isolates representation (its attention biased by Plücker-coordinate and Grassmann-window geometry rather than position alone), Shifu-LMR-Long isolates context (thousands of nucleotides), and LMR-bpRNA isolates the data (the v0 architecture trained on bpRNA-1m, not Shifu-Corpus). Full per-model specifications appear in Extended Data Table 2. Evaluating them properly, though, takes more than a single score, which the next section defines.

### Three axes: the Shifu Trifecta

Evaluating the Shifu-LMR family takes more than one number. The instability we opened with exposed a measurement gap that an audited dataset alone does not close: a single correctness score, on any benchmark, cannot say whether a model is broadly reliable or whether it knows when it is wrong. We therefore score every model on the Shifu-Corpus test set along three complementary axes: the Shifu Trifecta. The first is *correctness*: micro-F1, the base-pair accuracy used throughout the field. The second is *breadth*: the RNA Breadth Index (RBI), which asks whether a model stays usable across the hard, shifted parts of the data, combining how many sequences it can handle with how it does on its worst classes. The third is *awareness*: the RNA Uncertainty Index (RUI), which asks whether a model’s own confidence tracks its errors, so a user can see where a model is likely to struggle. A weather forecast is a useful analogy: correctness is how often it is right, breadth is whether it is right everywhere and not only in fair weather, and awareness is whether a stated “70% chance” can be believed. The exact formulas, which are defined on Shifu-Corpus’s evaluation units, are given in Methods (“Evaluation metric definitions”).

Awareness is operational: restricting a model to its most-confident predictions sharply raises accuracy when its confidence is trustworthy, but barely moves it when that confidence is uninformative, a behavior we make explicit on the Shifu-Corpus test split once the models are in hand.

Plotting breadth against awareness sorts models into four readable quadrants (Fig. 3a; the three axes shown together in Fig. 4). *Broad & Aware*, high on both, is the goal: broadly accurate and honest about uncertainty, where PriFold sits as the strongest external baseline. *Fragile & Aware* (low breadth, high awareness) describes a model whose point predictions are often inaccurate but whose confidence still flags them, so its outputs stay useful for ranking; BPfold is the clear case, at micro-F1 0.31 yet RUI 0.65. *Fragile & Blind* (low on both) is the regime to avoid, because the model fails *and* gives no signal that it is failing, as UFold does here. *Broad & Blind* is rare in our data. The three axes are genuinely independent (BPfold and UFold differ by 0.28 in RUI at similarly low micro-F1 (0.31 and 0.38), and BPfold reaches full coverage yet the lowest RBI among the full-coverage models), so no one axis can be read off another.

**Figure 3.**
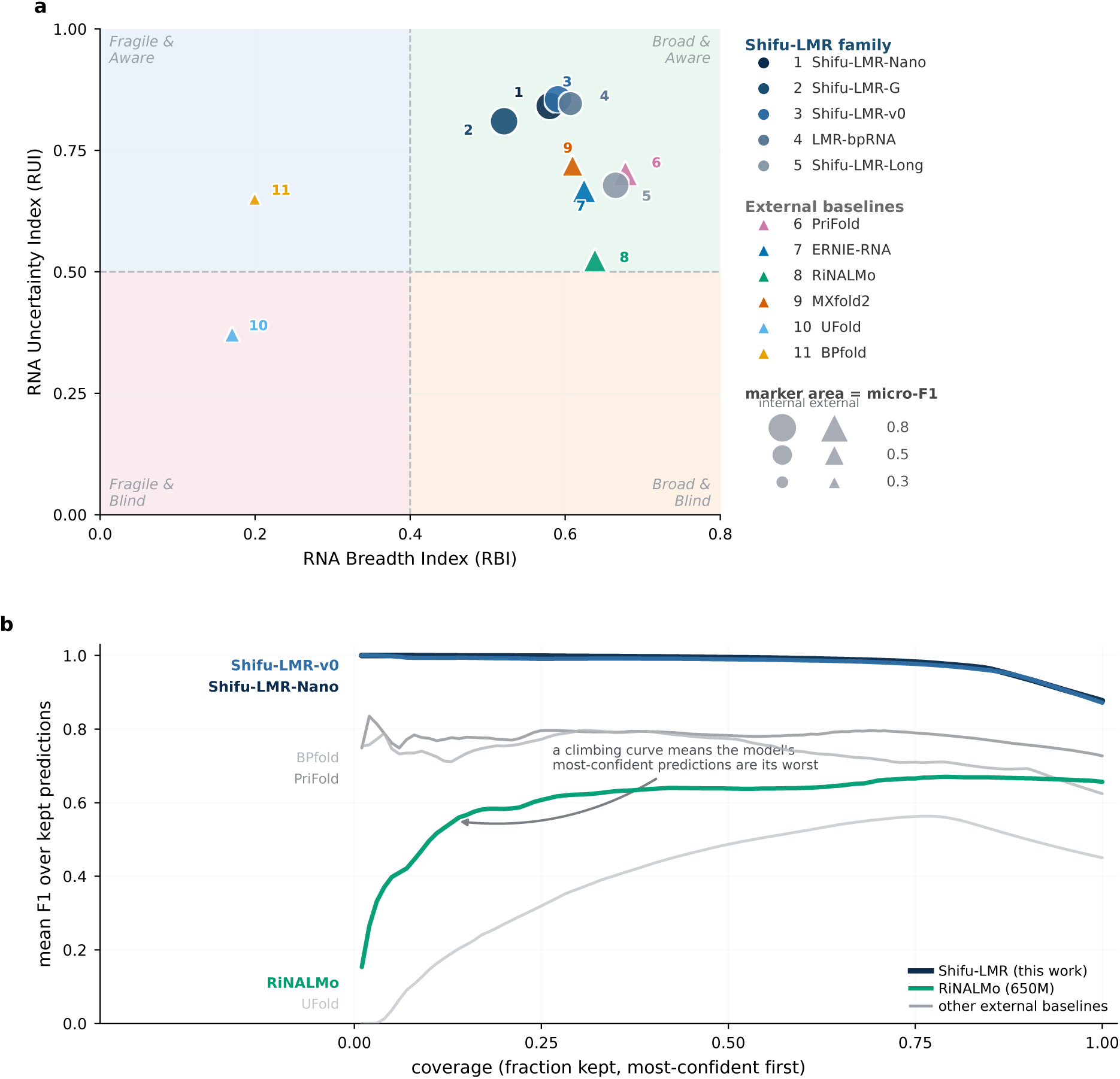
Breadth and awareness separate broad, aware models from benchmark specialists. **a**, the Shifu-Corpus test split (*n* =25413) on the breadth–awareness plane (RNA Breadth Index RBI, *x*; RNA Uncertainty Index RUI, *y*), marker area proportional to correctness (micro-F1) with quadrant cuts at RBI = 0.40, RUI = 0.50 (Methods). The Shifu-LMR models occupy *Broad & Aware*; among the externals PriFold anchors the same quadrant, BPfold is *Fragile & Aware* (low accuracy but informative uncertainty), and UFold is *Fragile & Blind*. **b**, selective trust: for each model, the mean per-sequence F1 over its most-confident fraction of predictions (lowest pairing entropy first), as coverage grows from the most-confident few to all sequences. A model whose confidence is trustworthy stays high when restricted to its confident predictions: the Shifu-LMR models do, whereas the 650-million-parameter RiNALMo’s most-confident calls are among its worst (the rising curve), so its confidence cannot gate its errors. These are the same per-unit curves that feed the ST-AUC term of RUI (Methods). Exact per-model values are in Table 1.

**Figure 4.**
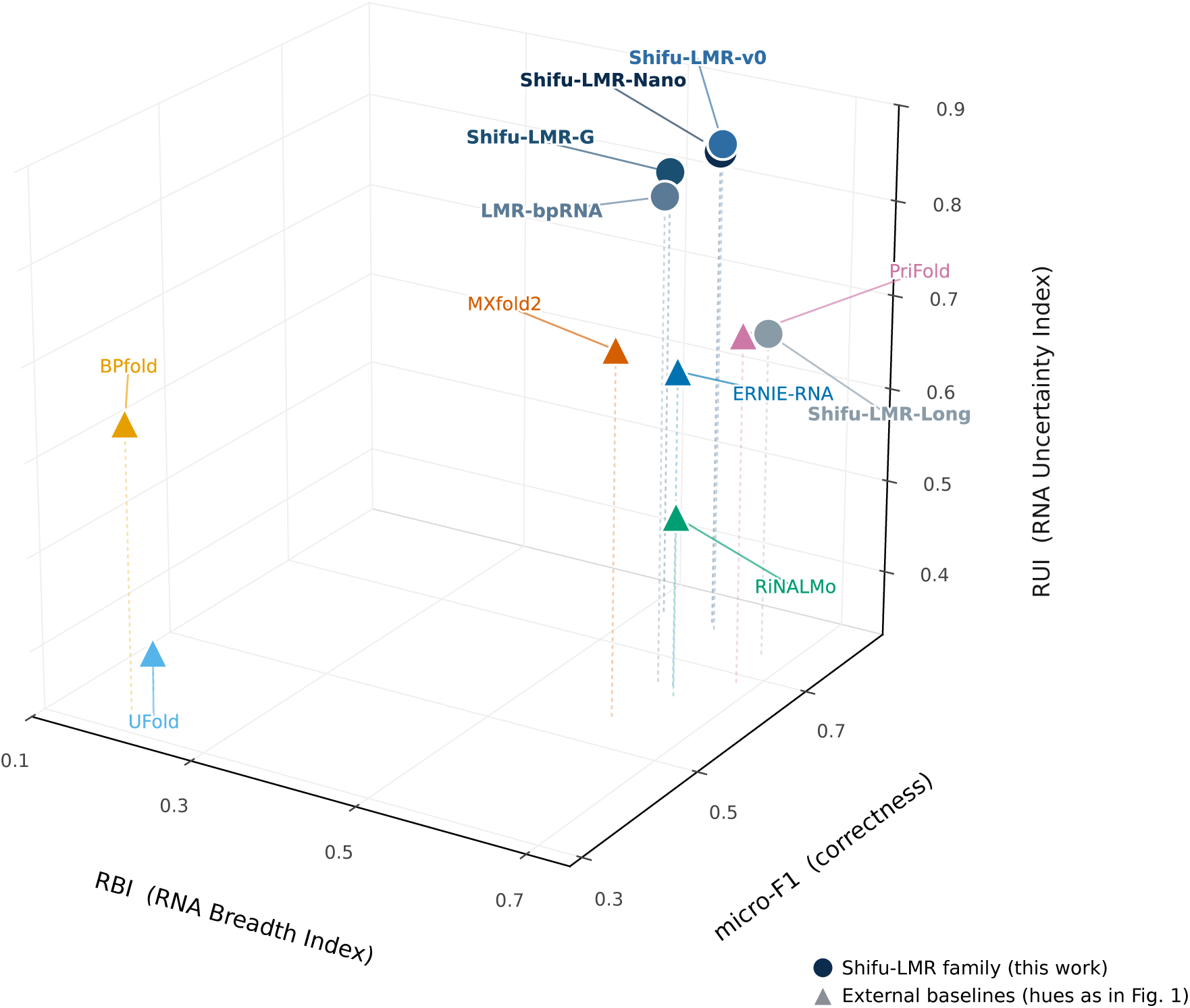
The Shifu Trifecta: three axes that one number cannot summarize. The Shifu-Corpus test split (*n* =25413) shown on all three axes at once: correctness (micro-F1), the RNA Breadth Index (RBI) and the RNA Uncertainty Index (RUI). The six external baselines are colored as in Fig. 1; the Shifu-LMR models of this work, introduced above, are drawn in a single muted navy-to-slate family so the figure reads as the evaluation framework rather than as a results comparison. The breadth–awareness quadrant map of the same models, with the cuts that define the four quadrants, is given in Fig. 3a; exact per-model values are in Table 1.

**Table 1.** The Shifu Trifecta on the Shifu-Corpus test split. (*n* =25413), for the internal Shifu-LMR family of this work (**a**) six external deep-learning baselines (**b**), and two non-deep-learning reference baselines (**c**). Best value per metric in bold; no single model leads on all three axes. Bootstrap 95% confidence intervals over test sequences place Shifu-LMR-Nano and Shifu-LMR-G within each other’s interval on micro-F1 (0.783– 0.793 vs. 0.782–0.792), both separable from Shifu-LMR-v0; RUI is not applicable (n/a) for EternaFold, whose single-structure output has no comparable per-position uncertainty. “Params” is the deployed (active) parameter count (internal: merged backbone + adapters + head; external: counted from each published checkpoint). “Ctx” is the maximum input length in nucleotides; BPfold (n/a) uses a dynamic-programming decoder with no fixed input window and is a six-model ensemble. The breadth index decomposes as 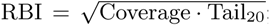; we show coverage (Cov, the fraction of sequences scored) and Tail20 (mean micro-F1 over the worst-performing 20% of evaluation units) so the index is transparent: BPfold, for example, scores every sequence yet collapses on its hardest fifth; per-model RBI and RUI decompositions are shown in Extended Data Fig. 2. micro-F1 (Eq. 1), RBI (Eq. 2) and RUI (Eq. 3) are defined in Methods; the validation split and per-corpus values are in Extended Data.

| Model | Params | Backbone | Training | Ctx | micro-F1 | RBI | Cov | Tail<sub>20</sub> | RUI |
| --- | --- | --- | --- | --- | --- | --- | --- | --- | --- |
| *(a) Internal: this work (Shifu-LMR family)* |  |  |  |  |  |  |  |  |  |
| Shifu-LMR-Nano | 65 M | LMR-Nano | Shifu-Corpus | 512 | **0.789** | 0.580 | 0.897 | 0.375 | 0.841 |
| Shifu-LMR-G | 88 M | LMR-G | Shifu-Corpus | 512 | 0.787 | 0.521 | 0.897 | 0.303 | 0.810 |
| Shifu-LMR-v0 | 291 M | LMR v0 | Shifu-Corpus | 512 | 0.776 | 0.591 | 0.897 | 0.389 | **0.855** |
| LMR-bpRNA | 291 M | LMR v0 | bpRNA-1m | 512 | 0.647 | 0.607 | 0.897 | 0.410 | 0.846 |
| Shifu-LMR-Long | 290 M | LMR-Long | Shifu-Corpus | 2,048 | 0.750 | 0.665 | 0.903 | 0.489 | 0.678 |
| *(b) External baselines* |  |  |  |  |  |  |  |  |  |
| PriFold | 182 M | T5-LM + RNAformer | bpRNA-1m | 488 | 0.683 | **0.678** | 0.894 | **0.514** | 0.703 |
| ERNIE-RNA | 87 M | RNA LM + head | bpRNA-1m | 1,024 | 0.648 | 0.624 | 0.902 | 0.432 | 0.668 |
| RiNALMo | 650 M | RNA LM (giant) | bpRNA-1m | 1,022 | 0.627 | 0.638 | 0.902 | 0.452 | 0.524 |
| MXfold2 | 0.8 M | thermodynamic + ML | TrainSetAB | 1,000 | 0.559 | 0.610 | 0.902 | 0.412 | 0.719 |
| UFold | 8.6 M | U-Net | bpRNA | 600 | 0.385 | 0.170 | 0.902 | 0.032 | 0.372 |
| BPfold | 48 M | transformer + DP | bpRNA | n/a | 0.314 | 0.199 | 1.000 | 0.040 | 0.650 |
| *(c) Non-deep-learning reference baselines* |  |  |  |  |  |  |  |  |  |
| EternaFold | small | CONTRAFold/Eterna CRF | Eterna | full | 0.531 | 0.634 | **1.000** | 0.402 | n/a |
| RNAfold | n/a | Turner NN (thermodynamic) | n/a | full | 0.460 | 0.608 | **1.000** | 0.369 | 0.664 |

Seen this way, the same three coordinates locate every model we evaluate, internal and external alike, on one map (Table 1); what the Shifu-LMR models in particular reveal we hold for the controlled experiment of the next section, where they are the subject rather than one more point on the plot. The take-home here is simply that three numbers, not one, are what let a reader tell a broadly trustworthy model from a benchmark specialist.

### Data and representation, not scale

Run through the Shifu Trifecta, the family answers the question it was built to pose: does data and representation, rather than parameter count, drive accuracy (Fig. 5)?

**Figure 5.**
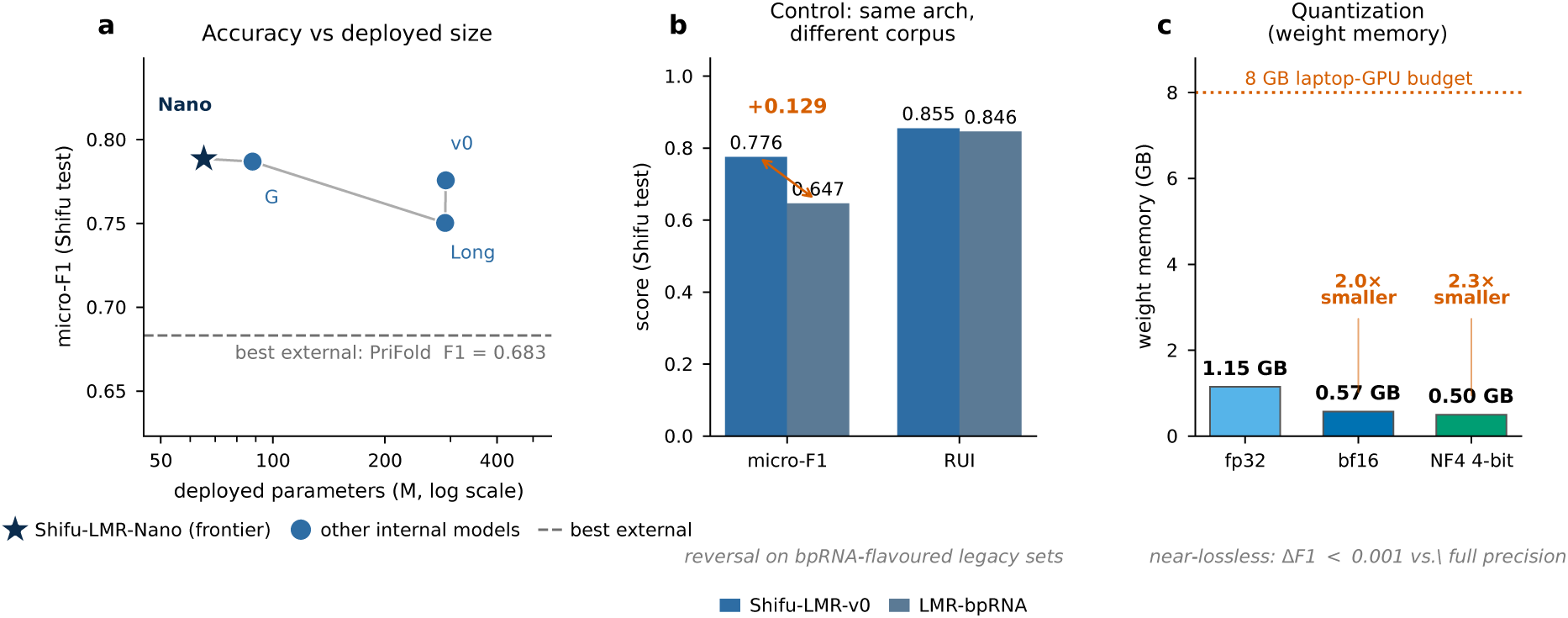
The Shifu-LMR family as a controlled experiment. **a**, correctness versus deployed parameter count on Shifu-Corpus test: Shifu-LMR-Nano (65 M) sits at the accuracy frontier, above the best external model (dashed line); **b**, the controlled corpus swap: the same architecture trained on Shifu-Corpus versus bpRNA-1m, a 0.13 micro-F1 gap at near-constant RUI; **c**, post-training quantization keeps the deployed model’s accuracy essentially unchanged (within 0.001 F1 of full precision; this is the checkpoint’s own inference accuracy, not the Shifu-Corpus-test micro-F1) while cutting weight memory from 1.15 GB to 0.50 GB (4-bit), within an 8 GB laptop-GPU budget. The masked-language-model pretraining, parameter-efficient adaptation, and staged-curriculum schematics for the family are shown in Supplementary Fig. S1.

The headline is Shifu-LMR-Nano, a 65-million-parameter model that reaches the highest correctness of any model we evaluate, internal or external (micro-F1 0.789 vs. 0.683 for the strongest external baseline; Table 1), together with strong awareness (RUI 0.841). Against RiNALMo, the largest external language model, the contrast is sharp: Shifu-LMR-Nano improves micro-F1 by 0.16 (0.789 vs. 0.627) with one-tenth the parameters (65 M vs. 650 M). The two axes beyond correctness tell complementary stories. Its breadth trails RiNALMo’s slightly (RBI 0.580 vs. 0.638), yet its uncertainty is far more useful (RUI 0.841 vs. 0.524), unsurprising because the external models were not built to report calibrated pairing confidence. It does so at a fraction of the size of the external language models and is light enough to run on a laptop GPU: evidence that, on a leakage-audited dataset, a compact model with a suitable training and representation strategy can match or exceed much larger ones on correctness.

Two classic, non-deep-learning tools anchor this comparison (Table 1). RNAfold and EternaFold fold every sequence (coverage 1.000) and outrank two published deep-learning models, UFold and BPfold, on correctness, while trailing the learned models overall. Their higher RBI reflects full length coverage rather than greater structural breadth: on the worst-performing fifth of units the compact learned models match them (Shifu-LMR-Nano Tail_20_ 0.375 vs. RNAfold 0.369).

To isolate the effect of the data alone, we hold the model architecture completely fixed and vary only the corpus it is trained on. The same 291-million-parameter model scores micro-F1 0.776 trained on Shifu-Corpus but only 0.647 trained on bpRNA-1m: a 0.13 gap from data alone, at essentially unchanged awareness (Fig. 5b). Tellingly, the relationship reverses on the bpRNA-flavored legacy benchmarks, where the bpRNA-trained model wins (TR0 0.84 vs. 0.77): each model is best on the distribution it was trained on (per-unit and per-family breakdown in Extended Data Fig. 4). This is the benchmark-bias effect of the first result, reproduced in-house under full control, with the internal models serving here as a controlled test of that effect rather than as the primary object of study.

Changing representation rather than data has a similar, independent effect. The geometric Shifu-LMR-G matches Shifu-LMR-Nano on correctness (0.787) at 88 million parameters, though with lower breadth: accurate but more fragile on the hardest classes (per-Rfam family performance is broken out in Extended Data Fig. 3). Extending context closes the breadth gap from the other side: a long-context variant (Shifu-LMR-Long) trades some peak correctness (0.750) for the highest breadth of any internal model (RBI 0.665), because it is not capped at short sequences. None of these gains came from adding parameters; each came from changing how data or structure is presented to a fixed-capacity model. One caveat applies throughout: because the internal models are trained on Shifu-Corpus, their edge over the externals measures the benefit of an audited, representative training set, the very effect we set out to isolate, while the leakage audit and seven held-out families keep the test genuinely held out.

Placed on the full Trifecta map, the Shifu-LMR models cluster in *Broad & Aware*: they lead the models evaluated here on correctness and on awareness while trailing the strongest external model on breadth, the one axis a short input window constrains (Fig. 3a). That awareness advantage is concrete. Restricted to their most-confident predictions the internal models stay accurate, whereas the 650-million-parameter RiNALMo’s most-confident calls are among its worst, so its confidence cannot gate its errors (Fig. 3b); this is the operational difference the RNA Uncertainty Index is built to capture.

Taken together, these results leave the field more than a leaderboard. They leave reusable practice: a curriculum that injects base-pairing and loop constraints so a model learns biological rules without new data; compact, biology-aware architectures that make scale a choice rather than a requirement; a leakage-safe way to build and split RNA data; and a three-axis vocabulary that turns “is this model good?” into the operational questions of whether it is broad enough and whether it knows when it is unsure. None of these is a final answer, but each is a step the community can build on.

## Discussion

The results above describe one problem seen from several sides. Across six external models and seven corpora the best model changes from benchmark to benchmark; within a benchmark, a single aggregate score hides the classes on which a model collapses; and on a leakage-audited dataset a 65-million-parameter model matches or beats far larger ones. Each view points to the same conclusion: RNA secondary-structure prediction is limited less by model scale than by how its data are curated, how RNA is represented, and how performance is measured. Much of the field’s recent optimism rests on aggregate micro-F1 climbing on a handful of bpRNA-derived benchmarks; our results suggest that part of that climb reflects overlap with training distributions rather than improved generalization, and that a single number can hide where a model is unsafe to use. Shifu is one response: a leakage-audited dataset, a measurement that reports breadth and awareness alongside correctness, and a family of controlled models in which better data and representation, rather than added parameters, are what moved accuracy.

This matters because of where the future of RNA model development lies. Structure predictions increasingly feed RNA therapeutics, target discovery, and synthetic biology, where a prediction must be accurate, honest about its uncertainty, and cheap enough to run at scale. A model that is accurate only on specialized hardware, or accurate but silent about when it is wrong, is hard to deploy responsibly. That a 65-million-parameter model runs on a laptop and still leads on correctness, and that the Shifu Trifecta reports when a prediction should be trusted, are steps toward predictions a practitioner can actually use. More broadly, the work treats machine learning as a deliberate scientific setup, much like framing a physics problem: the answer is only as good as the data, the constraints, and the objective one supplies, which is exactly why the data-and-representation view is the productive one. Shifu is aligned with, and significantly extends, the direction that RNASSTR began: RNASSTR showed that retraining fixed architectures on a homology-expanded, curated corpus improves secondary-structure prediction with no change to the model ^12^, establishing that better data, not new architecture, drives much of the gain. Shifu builds on that premise but goes further, pairing a leakage-audited corpus with an integrated three-axis evaluation and a family of controlled language models, so that the data, the measurement, and the model are addressed together. The RNASSTR labels, being covariation-derived, also carry caveats that an audited, mixed-fidelity corpus is designed to expose.

To make the dataset and evaluation usable by others, we release the full Shifu-Corpus audit suite up front (the splits, per-pair leakage and near-duplicate certificates, per-source label coverage, and byte-level manifest hashes) so that any group can verify the exact data a result was computed on rather than reconstruct it. This speaks to a recognized pain point: benchmarking RNA structure prediction is sensitive to dataset choice and metric definition, and the community has repeatedly cautioned that headline accuracies are hard to compare across studies ^5,8^. By pinning the data, the splits, and three orthogonal metrics, Shifu turns that comparison from an act of faith into one that can be audited.

The approach has clear limits, which we state plainly. The Breadth and Uncertainty indices are defined on Shifu-Corpus’s own evaluation units and length bands; their values are native to this dataset and are not intended as universal metrics or as a theory of calibration, and the quadrant cuts are interpretable anchors rather than fixed thresholds. The comparison that favors our compact models is bounded in the same way: they are trained on Shifu-Corpus, so their edge over the externals reflects training on a representative, leak-free set, the effect we set out to measure, which the same-architecture corpus swap isolates directly. Our standard models also cap input at 512 nucleotides and so trail on breadth, a gap the long-context variant only begins to close; and Shifu-Corpus inherits the compositional skew of public data, which the splitter mitigates but does not erase. Finally, the Trifecta scores exact base pairs and does not credit single-nucleotide register shifts; this is a deliberate scope choice, and an exploratory one-nucleotide-tolerant recount (Supplementary Table S3) confirms it does not change the ranking or the conclusions.

For users, the three axes double as a decision guide. A pipeline that needs dependable predictions across diverse RNAs should prefer a *Broad & Aware* model; one that ranks many candidates and validates the top hits downstream can still exploit a *Fragile & Aware* model by trusting its confidence; and a *Fragile & Blind* model warrants caution, because it gives no signal of its own failures. None of this is the last word. We see it as one step toward treating RNA structure prediction as a discipline in its own right, in which data curation, model representation, and evaluation are designed together. The same pattern (audit the data, stratify the evaluation, and report more than one number) should carry to pseudoknot-aware and, eventually, 3D RNA modeling ^24^. These concerns are, if anything, sharper for threedimensional structure. RNA3DB^15^ applies a comparable leakage-aware, clan-based split with careful covariance-model cutoffs to experimentally determined 3D structures, but that pool is far smaller and more skewed than the 254123 secondary-structure sequences assembled here: Rfam lists on the order of 150 families with experimental 3D structures, against the much larger family space represented at the secondary-structure level. The data-and-representation limits documented here for 2D therefore bound 3D modeling at least as tightly, and the same audit-first discipline will be needed there. We release the dataset, the evaluation code, the per-unit measurements, and the model backbones so that others can adopt, audit, and build on all three contributions, and so the field can tell genuine progress from benchmark artifacts as RNA structure prediction matures.

## Methods

### Dataset construction and deduplication

Complete per-source and per-split curation statistics, the full audit manifest, and the tiered-labeling breakdown are given in Supplementary Note 1 and Supplementary Table S1; we summarize the construction here.

Shifu-Corpus ingests records from six public RNA secondary-structure sources, each chosen for the slice of the labeling-density and structural-fidelity range it covers: bpRNA-1m at 90% identity clustering ^9^ for curated dot-bracket supervision; ArchiveII ^10^ for hand-curated, family-stratified records spanning canonical RNA classes (5S, 16S, 23S rRNA; group I/II introns; RNase P; SRP; tRNA; tmRNA; telomerase); RNAStrAlign^11^ for alignment-derived structures with explicit family and subfamily labels across ten families; RNASSTR^12^ for a large bulk corpus that provides scale (no pre-existing family annotation; family labels inferred post-hoc by the Infernal pipeline below); DSSR-derived structures at 90% identity^13^ for physics-grounded 3D-to-2D projections of riboswitches, ribozymes, and other classes that benefit from coordinate-derived ground truth; and RNAsolo (the cleaned PDB-derived RNA structure repository of the RNApolis platform), which contributes hand-curated PDB chains carrying pdb id, chain id, method, resolution, and release date fields that allow date- or resolution-stratified evaluation. Rfam family annotations from the Rfam clan-input file^25^ are used as splitting metadata only; Rfam itself is not ingested as a record source.

Per-record processing harmonizes the six sources to a common schema. Sequences shorter than 20 nt or longer than 8192 nt are excluded from the active set; the lower bound removes truncated fragments that have insufficient structure to evaluate, and the upper bound covers the longest supported input across the inference pipelines we evaluated. Each ingestor parses the source’s dot-bracket notation, including multilevel bracket–letter encodings of pseudoknots (⟨( ) [ ] { } ¡ ¿ Aa-Zz⟩); the parsed pair list is split into a nested set and a pseudoknot set, with the pseudoknot set preserved in a has pk raw flag and the canonical no-pseudoknot dotbracket retained as the prediction target. Because the scored reference is pseudoknot-stripped, a predicted pair that correctly recovers a pseudoknotted contact is counted as a false positive, so micro-F1 under-credits models that predict pseudoknots; pseudoknotted pairs are 2.4% of test pairs, and this penalty is not uniform across models because the nested Watson-Crick and wobble decoders used by the internal models cannot emit pseudoknots and so cannot incur it. We release the raw pair lists and the has pk raw flag for pseudoknot-aware re-scoring. We adopt this convention because most current models predict nested structures only, and silent loss of pseudoknot information would make cross-model comparison opaque. Pairs are normalized to DSSR canonical types (A–U, G–C, G–U); non-canonical pairs are removed from the supervision target but preserved in the raw pair list for downstream inspection.

Each record receives a deterministic 24-character UID derived from the SHA-256 of the tuple (source, source id, sequence, first-20-pairs-json). The UID is the join key for downstream operations (Infernal evidence merge, evaluation slice manifests, eval-export CSVs) and lets external auditors verify that an evaluation run consumed the same records the manuscript reports. UID uniqueness is asserted before any downstream phase; on collision the build aborts. Exact deduplication retains the first occurrence of each distinct sequence, identified by the SHA-256 fingerprint of the normalized sequence. Of 310034 ingested records, 55641 are removed as exact duplicates, yielding 254393 unique sequences; a subsequent 20–8192 nt length filter removes 270 more, leaving 254123. The ingestion order is bpRNA first; when a sequence appears in both bpRNA and another source, the bpRNA copy is retained. We adopt this convention because bpRNA carries the most uniformly curated supervision among the six sources, and because retaining the copy with the highest-quality structure annotation reduces the risk of a downstream model being trained against a less reliable label.

Near-duplicate clustering operates on the deduplicated records to prevent cluster-level leakage that exact dedup cannot detect. MinHash signatures^26^ are computed over the set of 6-mers of each sequence with 128 permutations, and records are merged into the same cluster when their estimated Jaccard similarity meets or exceeds 0.90. We chose MinHash because exact pairwise Jaccard would scale quadratically with dataset size (infeasible at 254 k records), and because Jaccard over *k*-mers is a standard proxy for sequence similarity in bioinformatics ^27^. The 6-mer choice balances locality (smaller *k* oversmooths) and discrimination (larger *k* under-smooths); 128 permutations gives a Jaccard estimate with standard error ≈ 0.044, sufficient for the 0.90 threshold. The Jaccard threshold of 0.90 is a deliberate choice: it removes near-identical sequences that would otherwise leak across splits, while preserving distinct sequences that share family-level homology. Cluster identifiers are persisted per record and become the join key for the leakage audit.

Family and clan labels are produced by a three-tiered, evidence-tracked procedure. A first heuristic pass maps known family names from ArchiveII and RNAStrAlign to Rfam accessions through a curated table (e.g., trna → RF00005; 16s rrna → RF00177/RF01959/RF02542). A second pass runs Infernal cmscan ^28^ against the Rfam covariance-model database with --cut ga --rfam --nohmmonly --fmt 2 and the clan-input file. The --cut ga flag applies each covariance model’s curated gathering threshold, the canonical Rfam-team criterion for “this is a genuine hit”. We classify each record’s labels into three tiers: *gold* requires the inclusion mark ! (above the gathering threshold); *pred* accepts the best hit regardless of confidence; and *active*, the tier used in evaluation, takes the gold label when available, falls back to a high-confidence prediction (bits ≥ 25) when available, falls back to the heuristic name-mapping when neither Infernal hit type is available, and finally to *unlabeled* otherwise. Of the 254123 active records, 193650 (76.2%) carry a gold label and 223871 (88.1%) carry an active label. The released dataset carries all three tiers per record together with the underlying evidence (rfam topk, rfam best per clan, rfam pred evalue/bits/inc), so any future tier reclassification is auditable in place. We adopted the tiered design because a single Infernal-only label policy would discard roughly 11% of the dataset for which the heuristic mapping is the strongest available evidence, while an Infernal-or-heuristic policy without explicit tier tracking would conflate evidence sources of differing quality.

### Family-aware splitting and leakage audit

Sequences are split into train/validation/test partitions with target ratio 80/10/10 by a greedy stratified group-split. Each record receives a group key by the priority order Rfam clan → Rfam family → non-provenance raw family label → unique singleton when no biological grouping is available. We adopt this priority because clan is the broadest biological grouping and therefore the most leakage-relevant: two records sharing a clan share enough structural homology that allowing one into train and the other into test risks training-set contamination of the test-set evaluation. Records with no biological annotation are assigned unique singletons rather than merged into a single *unknown* mega-group, because the latter would force all unannotated records into one split and cause large, biologically heterogeneous transfers between splits.

These per-split counts should not be read as the corpus’s family diversity. Shifu-Corpus is predominantly unlabeled at the Rfam level (records without a confident Rfam assignment, shown pooled as *unknown*, are 88.8% of train and 86.2% of test), so only a small number of Rfam families are large enough to appear as distinct groups per split (four, two, and eight for train, validation, and test, each counting the pooled unknown group). Rfam accessions are used to define the family-disjoint held-out test families (Supplementary Table S2), not to enumerate the corpus, which is a secondary-structure resource drawn from six databases rather than a collection of experimentally determined 3D structures.

The greedy assignment processes super-groups largest-first (most-constrained-first) and scores each candidate split on a weighted sum of (i) the L1 deviation between the trial split’s realized ratio and the target 80/10/10, and (ii) the L1 deviation between the trial split’s source distribution and the global source distribution, with a 2× weight on source-mix divergence. We adopted the 2× source-mix weight because without it the dominant RNASSTR source (76.9% of the dataset) would saturate any one split; pilot runs without this penalty produced splits with *>* 99% RNASSTR in val/test, which would make cross-source generalization unmeasurable. Super-groups exceeding max(500, 2% of dataset) records are shattered before assignment, preserving near-duplicate cluster integrity within each chunk; this prevents pathological mega-clans (notably SSU rRNA at *>* 150000 records) from single-handedly determining the split. Length stratification uses ten quantile-based bin edges computed globally and persisted in audits/split summary.json for reproducibility.

The seven Rfam families absent from train and validation (RF00028, RF00017, RF00009, RF00023, RF02541, RF00029, RF00024) are an emergent property of the seed-42 stratified assignment, not an *a priori* reservation. The slice builder post-certifies the family-heldout invariant set(test family heldout.families) ∩train families = ∅ ø and aborts the build on violation. A different seed could place different families in test; we report seed 42 as the canonical release and the audit as the guarantee. Groups proposing additional Shifu-Corpus releases under different seeds are encouraged to publish their audit alongside, so cross-release comparisons remain meaningful.

The leakage audit runs after splitting. Exact-identity leakage is tested by checking whether any sequence appears verbatim in two splits; cluster leakage is tested by checking whether any MinHash cluster spans two splits. All six pairwise checks (exact and cluster, for each of train↔val, train↔test, and val↔test) return zero in the v13 release. The audit JSON is included in the release and the build aborts on any non-zero count. We treat this audit as an integrity proof rather than a quality measure: it does not guarantee absence of homology, only absence of near-duplicate sequence identity at the chosen Jaccard threshold.

### Evaluation units

Each model’s predictions are partitioned into evaluation units defined by (source, length-bin), with length bins short (≤ 200 nt), medium (201–600 nt), and long (*>* 600 nt). A unit with fewer than 30 sequences is merged into the next larger length-bin within the same source; if no bin in the source has at least 30 sequences, the source is collapsed into a single *all* unit. The 30-sequence threshold reflects the smallest unit at which Tail_20_ (defined below) is robust to single-sequence outliers; smaller units would let one bad prediction dominate the unit-level score. This merging rule yields 11 evaluation units on the Shifu-Corpus test split for the Trifecta computations (Table 1, Fig. 3); the high-resolution per-unit view in Extended Data Fig. 1 retains all 18 pre-merge cells for visibility.

### Evaluation metric definitions

Micro-F1 is the base-pair-level F1:

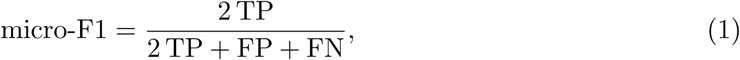

where TP (true positives), FP (false positives) and FN (false negatives) are summed over all sequences in the evaluation set. A predicted pair (*i, j*) is a true positive when the ground-truth structure contains it, a false positive when it does not, and a ground-truth pair the model misses is a false negative. We report micro-averaged rather than macro-averaged F1 because RBI carries the unit-level treatment separately; macro-averaging at the F1 axis would double-count the inhomogeneity that RBI is designed to expose.

The RNA Breadth Index (RBI) measures breadth of competence across hard, shifted evaluation units:

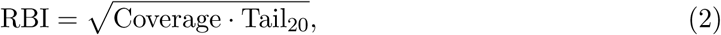

where Coverage = *n*_proc_*/n*_total_ is the fraction of split sequences for which the model returned predictions (with capacity-exceeding sequences receiving a discounted score; below), and 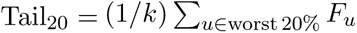 is the unweighted mean micro-F1 over the worst-performing 20% of post-merge evaluation units (with ties broken toward smaller units). We use the geometric mean rather than the arithmetic mean so that neither factor alone produces a high RBI; in particular, RBI = 0 when either coverage or tail performance is zero. This penalizes models whose narrow specialization gives them strong head-of-distribution F1 but no predictions on out-of-distribution units, and equally penalizes broadly mediocre models whose tail performance is uninformative.

The RNA Uncertainty Index (RUI) measures whether a model’s per-residue uncertainty estimates rank its errors usefully:

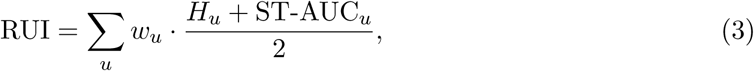

where *H_u_* = (1 + *ρ_u_*)*/*^2^ maps the unit-level Spearman rank correlation *ρ_u_* = *ρ*(entropy, error) ^29^ into [0, 1]; 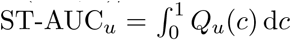 d*c* is the area under the selective-trust curve ^30,31^, with *Q_u_*(*c*) the mean micro-F1 over the lowest-entropy fraction *c* of sequences in unit *u*; and *w_u_* is proportional to unit sequence count, normalized so that 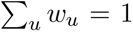. We use Spearman rather than Pearson rank correlation because the metric should not require entropy-scale calibration; only the ordering need be informative. We use (*H_u_* + ST-AUC*_u_*)*/*2 because the two components measure empirically distinct uses of uncertainty (rank correlation between entropy and error vs. the thresholded-coverage F1 curve), and we did not find evidence to weight one over the other in our six-model data.

Entropy quantifies how much a probability distribution commits to an outcome, the standard information-theoretic measure of uncertainty ^32,33^, and the view of that uncertainty as an accountable physical quantity has deep roots in the physics of information ^34,35^. RUI makes this operational: it asks whether a model’s per-residue pairing entropy ranks its own errors, so that a pipeline can gate low-confidence predictions ^36^. We position RUI as an operational measure of uncertainty usefulness rather than as a universal calibration measure. The distinction matters: RUI answers the operational question *can a downstream pipeline trust this model’s confidence to gate predictions on Shifu-Corpus’s units*, not the broader decision-theoretic question of whether a model is well-calibrated across all possible inputs and decision rules ^49^. We caution against extrapolating RUI to settings outside the Shifu-Corpuson which it is defined.

Sequences exceeding a model’s declared maximum input length are not silently dropped from the evaluation. Each receives a capacity-discounted score *F_i_*= *F*_base_ · (*l*_max_*/L_i_*)*^β^*, where *F*_base_ is the mean micro-F1 of the top 10% of in-range sequences by length, *L_i_* is the sequence length, *l*_max_ is the model’s declared maximum input length, and *β* = 1.0 throughout Table 1. Without this discount, a model could earn an artificially high RBI by refusing to score difficult sequences; with the discount, the model is held responsible for sequences it cannot process. We chose *β* = 1.0 as a conservative default; alternative values are reported in the per-unit data release for sensitivity analysis.

Quadrant cuts are RBI = 0.40 and RUI = 0.50. We chose 0.40 for RBI as the geometric mean of 80% coverage with a 20% tail-floor, an operational threshold that separates broadly usable from narrowly usable models in our six-model field. We chose 0.50 for RUI as an operational cut anchored on the ranking term: *ρ_u_* = 0 (an uninformative error ranking) corresponds to *H_u_* = 0.5, so RUI = 0.5 is the no-information level for that term. The selective-trust term has no fixed no-information value (its baseline is a model’s own accuracy), so we treat RUI = 0.5 as a practical rather than exact boundary between informative and uninformative uncertainty. Other cuts are defensible; we treat these defaults as operationally interpretable rather than as universally correct.

### Model evaluation protocol

Six external models were evaluated under the Shifu Trifecta protocol: ERNIE-RNA ^18^, BPfold ^23^, RiNALMo ^20^, UFold ^22^, PriFold ^19^, and the thermodynamic-plus-machine-learning hybrid MX- fold2 ^21^. Each was selected for the architectural and methodological niche it occupies: three pretrained RNA language models with different pretraining corpora (ERNIE-RNA, RiNALMo, PriFold), one transformer with a classical decoder (BPfold), one U-Net-style convolutional model (UFold), and one thermodynamic+ML hybrid (MXfold2). Together they let the six-model field span the architectural classes currently active in the literature. Each model was evaluated at native precision with the authors’ recommended decoding parameters; what we modify is the export surface, not the prediction itself.

The Trifecta requires a per-sequence symmetric pairing-probability tensor *P* ∈ [0, 1]*^L×L^* so that per-residue entropy can be computed and the selective-trust curve built; few published inference pipelines return this tensor by default. For each model we therefore added a thin evaluation wrapper that (i) runs the published forward pass with the published weights and tokenizer, (ii) intercepts the pre-symmetrization pairing logits at the last layer before the model’s built-in decoder (or at the pairing head directly where one is exposed), (iii) applies the model’s own symmetrization rule (transposition with a max or mean reduction, following the published code), and (iv) passes the thresholded structure through the same base-pair evaluator used to compute micro-F1. The decoded structure used to compute F1 is identical to what the model’s public release produces; the added *P* tensor is used only for *H_u_* and ST-AUC*_u_*. Per-residue entropy is 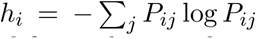 ^32,37^ with the convention 0 log 0 = 0, and the per-sequence entropy used for ranking is the mean of *h_i_* over residues. RiNALMo’s tokenizer rejects a small number of sequences containing non-canonical residues; these are excluded from its evaluation units, with the exclusion absorbed into the coverage term of RBI rather than ignored.

Two practical notes apply to the model field. PriFold’s published release contained an off-by-one tokenization slice that we identified during evaluation; we corrected this in our wrapper and report numbers consistent with the architecture’s expected behavior. We initially planned to include MetaFold-RNA but its published inference pipeline requires a four-tool ensemble prior (RNAfold, LinearFold-C, LinearFold-V, RNAstructure) on ∼175000 sequence-runs that we could not reproduce without introducing a prior-construction bias; we substituted MXfold2, whose self-contained inference pipeline and architectural distinctness from the four pure-deeplearning models make it a better single comparator across architectural classes. Both choices are documented for reproducibility.

### Foundational RNA language models

The internal models are built on a family of transformer encoders ^38^ pretrained on unaligned RNA sequences from RNAcentral ^39^ with a masked-language-modeling objective ^40^, then reused across structural tasks. We pretrain three backbone sizes spanning a deployment range: a 289-million-parameter model (LMR v0; *d*_model_ = 1024, 24 layers, rotary position encoding ^41^, SwiGLU feed-forward ^42^), a 64-million-parameter model (LMR-Nano; *d*_model_ = 576, 12 layers) that is the basis of Shifu-LMR-Nano, and a geometric variant (LMR-G; 86 or 228 M parameters), introduced here, whose self-attention is biased by Plücker-coordinate and Grassmann-window features inspired by Grassmann-Plücker sequence flows ^43^, so that it is sensitive to the relative geometry of nucleotides rather than position alone. A long-context backbone (LMR-Long) extends the receptive field to 4096 nucleotides through hybrid sliding-window and full attention. Pretraining is self-supervised and uses no structural labels; these backbones are the basis on which structure is subsequently learned.

### Structural fine-tuning by curriculum

Because a nucleotide sequence underdetermines its structure, we do not fine-tune the backbone end-to-end on pairing labels directly. We freeze the pretrained backbone and attach a small set of trainable components: low-rank adapters (LoRA)^44^, with weight-decomposed adapters (DoRA) ^45^ for the geometric backbone, and an axial pairing head that emits a symmetric pairing-probability tensor, which a nested dynamic-programming decoder with Watson–Crick and wobble constraints converts into a structure. The pairing head is trained with a focal objective ^46^ to counter the extreme sparsity of true base pairs. Supervision is delivered as a staged *curriculum*, a recipe rather than a fixed schedule, that introduces biological constraints progressively rather than all at once: a soft-label warm-up on Rfam secondary-structure-conserved alignments, pairing initialization, joint training gated on a pairing-F1 threshold, and hard-negative mining. Not every model uses every stage (Extended Data Table 2), and the schedule can be shortened or extended. The curriculum presents the same data in order of increasing structural context (canonical pairing rules, then loop-length and one-partner-per-base constraints, then global consistency) so the model acquires the rules that govern folding without additional data. This is the operational form of the representation argument of the Introduction: we add context, not volume.

### Hardware and compute

All training and evaluation were performed on a single workstation with two AMD EPYC 9354 32-core processors (64 physical / 128 logical cores), 1.5 TB of system memory, and eight NVIDIA RTX 5000 Ada Generation GPUs (32 GB each; 256 GB aggregate), hosted by the University of Houston Enterprise Systems Facilities. The structure-prediction models reported here were fine-tuned on a single GPU; backbone pretraining used one or more of these GPUs. To substantiate the “runs on a laptop” claim, Shifu-LMR-Nano inference was additionally validated on a consumer laptop (Intel Core Ultra 9 with an NVIDIA RTX 5050 Laptop GPU, 8 GB).

### Quantization

The released models support post-training quantization so they fit and run within a small memory budget without retraining. Casting weights to half precision (float16/bfloat16) ^47^ is essentially lossless (base-pair F1 changes by less than 0.001) and halves weight memory, while 4-bit weight quantization of the frozen backbone (NF4)^48^ cuts weight memory by more than half (for the long-context model, from 1.15 GB at full precision to 0.50 GB at 4-bit) at near-unchanged accuracy (micro-F1 ≈0.96) and a modest per-sequence slowdown. The geometric backbone is quantized at its linear layers only, leaving its Plücker-coordinate attention in full precision. Together these let the 65-million-parameter Shifu-LMR-Nano, and even the 291- million-parameter pairing models, serve within the 8 GB budget of a laptop GPU.

### Training and curriculum

All structural models are trained on a single GPU with a frozen backbone, effective batch size 32, mixed bfloat16 precision, and a maximum input length of 512 nucleotides. The structural variants reported here were trained on the Shifu-Corpus training split, with the sole exception of LMR-bpRNA, which is identical in architecture to Shifu-LMR-v0 but trained on bpRNA-1m at 90% identity for the controlled corpus comparison in the Results. Backbone pretraining dominates training cost (for example, the LMR-Nano backbone trained for roughly 145 GPU-hours); structural fine-tuning adds a smaller increment, and the bpRNA-trained pairing model completed in under 40 hours. All parameter counts are reported at full precision. Per-model parameter counts, backbone and adapter configurations, and curriculum schedules accompany the code release as a general recipe; the exact per-model hyperparameters of the withheld models are available to editors and referees on request, and the staged-curriculum details and the entropy and metric derivations are given in Supplementary Information.

### Statistics and reproducibility

All Trifecta values reported in Table 1 and in Fig. 3 are deterministic summaries of the evaluation set: micro-F1 is computed from the full per-sequence TP/FP/FN tally; RBI and RUI are unit-stratified summaries with unit weights *w_u_* proportional to unit sequence count. Beyond a per-sequence bootstrap of pooled micro-F1 (reported as the 95% confidence intervals in Table 1), no additional bootstrapping or error-bar treatment is applied at the split level; the per-unit values are released so that any user wishing to bootstrap unit boundaries or re-weight units can do so. Aside from those micro-F1 intervals, the Shifu Trifecta is a reporting protocol rather than a statistical test, and we make no formal claim of pairwise significance on the breadth and awareness axes. The full build is deterministic from a single seed (default 42); the MinHash permutation family, the super-group shuffle, and the global RNG all consume from this seed. Manifest hashes for each split file are published so that any reconstruction can be checked byte-for-byte.

### LLM assistance disclosure

Large language model tooling was used during manuscript preparation for editing assistance, prose suggestion within author-written scaffolds, and consistency checking against the audit JSON files. All scientific decisions, the full numerical content, the metric definitions, the figure designs, and the final wording were author-determined and author-verified against the released audit artifacts. No LLM is listed as an author and no LLM was assigned a CRediT role.

## Data availability

The full Shifu-Corpus, including train, validation, and test splits, is released on HuggingFace at https://huggingface.co/datasets/GaboG7/SHIFU-Corpus and will be deposited under a permanent archival DOI on publication. Each released record carries the normalized RNA sequence; the raw and pseudoknot-stripped dot-bracket structures; the parsed pair lists; per-record statistics (length, GC, pair density, canonical-pair fraction, `has pk raw`); the source dataset and source identifier; the three tiered Rfam family/clan labels (`rfam pred *`, `rfam gold *`, `rfam active *`) with the active-policy column and perrecord confidence scalar (`rfam active conf`); and, for records where Infernal returned hits, the full evidence set (`rfam best overall`, `rfam topk`, `rfam best per clan`, `rfam pred evalue/bits/inc`, `rfam n hits`). PDB-derived records additionally carry `pdb id`, `chain id`, `method`, `resolution`, and `release date`. Four pre-computed evaluation slices (`test standard`, `test family heldout`, `test clan heldout`, `test unlabeled ood`) are released as UID-only manifests. The audit suite (`global summary.json`, `split summary.json`, `dedup report.json`, `leakage report.json`, `label coverage.json`, `tiered label summary.json`, `edge case counters.json`, `parity diagnostics.json`, `manifest hashes.json`) is released alongside the dataset.

## Code availability

All dataset-construction code, the Infernal annotation driver, evaluation scripts, metric implementations, and figure-generation code are released under an open license at https://github.com/Vicens-Lab/LMR, with a tagged release matching this manuscript and a permanent archival DOI on publication.

## Models availability

The foundational model backbones (LMR-v0, LMR-G, LMR-Nano, and LMR-Long) are released on HuggingFace at https://huggingface.co/GaboG7/LMR-v0, https://huggingface.co/GaboG7/LMR-G, https://huggingface.co/GaboG7/LMR-mini, and https://huggingface.co/GaboG7/LMR-Long, respectively. The structure-prediction models (the Shifu-LMR family reported in Table 1) are reproducible from the released data, code, and training configurations; trained weights are available from the corresponding author on request.

## Protocol availability

A protocols.io deposition describing the end-to-end Shifu-Corpus evaluation procedure (download, infer, compute Trifecta, generate figures) is released alongside the code repository.

## Acknowledgements

We gratefully acknowledge Anton I. Petrov and Nivedita Dutta for careful reading of the manuscript; John Gillet and the UIT at the Enterprise Systems Facilities for hosting our workstation; and University of Houston start-up funds (to Q.V.).

## Author contributions

G.G. and Q.V. conceived the project and designed the research. G.G. developed the computational framework, performed the data analysis, and generated the figures. Q.V. supervised the study and provided project resources. G.G. wrote the initial draft of the manuscript, and both authors contributed to the final editing and revision of the text.

