## Extended data for "Shifu: an integrated framework for deep learning of RNA secondary structure"

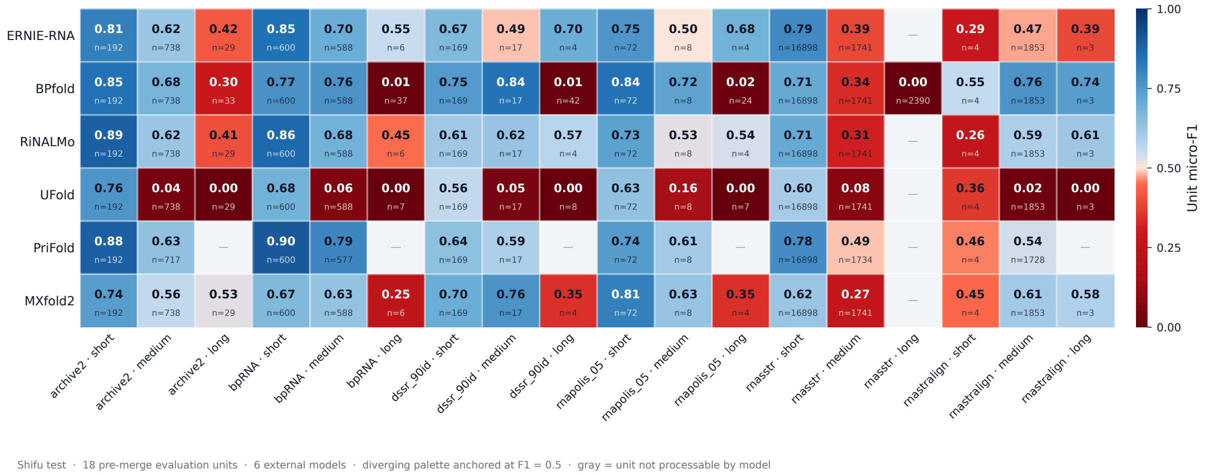


**a.** Shifu-Corpus test split.

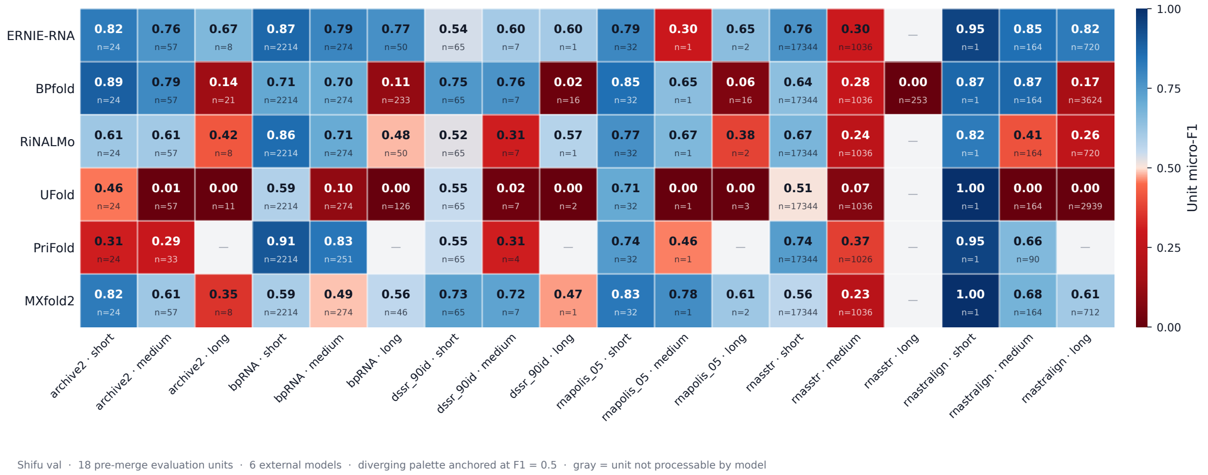


**b.** Shifu-Corpus validation split.

**Extended Data Fig. 1. Within-benchmark fragility: aggregate metrics hide unit-level failure.** Per-unit micro-F1 across (source × length) units on the Shifu-Corpus test (**a**) and validation (**b**) splits; gray cells mark units a model cannot process (sequences beyond its declared length limit). Every external model collapses to near-zero accuracy on at least one unit, typically the longest sequences, and the failing units differ across models, so two models with the same aggregate micro-F1 can fail in entirely different places. Sample sizes: Shifu-Corpus test $n$ =25413, validation $n$ =25412.

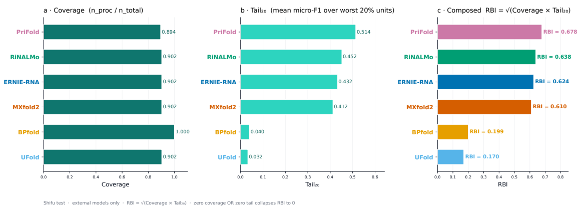

**a.** RBI components, test.

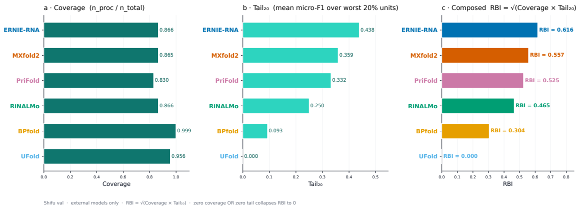

**b.** RBI components, val.

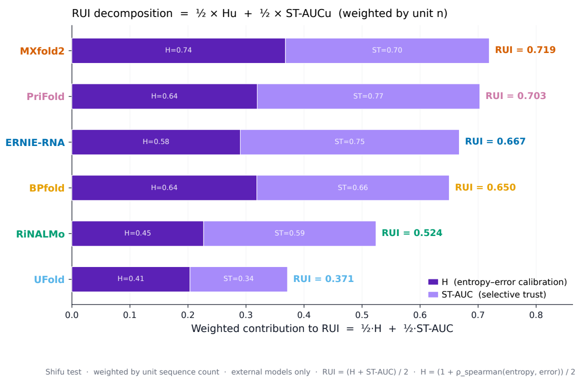


**c.** RUI components, test.

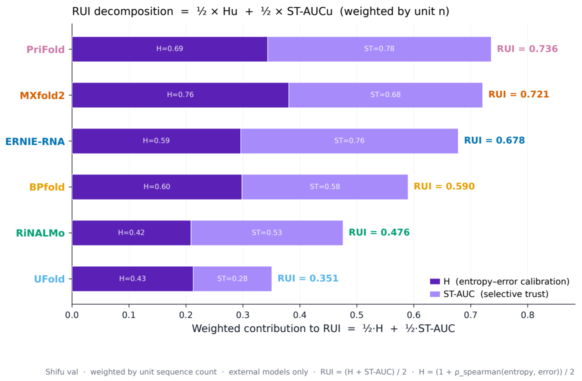


**d.** RUI components, val.

**Extended Data Fig. 2. RBI and RUI decomposed into their components. a, b**, Coverage, $\text{Tail}_{20}$, and the geometric-mean composed RBI $= \sqrt{\text{Coverage} \cdot \text{Tail}_{20}}$ on test (a) and validation (b). On test, BPfold reports coverage 1.000 but $\text{Tail}_{20} = 0.040$, so RBI collapses to 0.199; UFold's tail is 0.032, driving RBI to 0.170; PriFold, ERNIE-RNA, RiNALMo, and MXfold2 earn both factors. **c, d**, Weighted contributions of $H$ (entropy–error rank correlation, deep violet) and ST-AUC (selective-trust area, light violet) to composite RUI $= (H + \text{ST-AUC})/2$ on test (c) and val (d). Because $H$ is rescaled to $[0, 1]$ before weighting, BPfold's low micro-F1 does not preclude a strong $H$ contribution; on test, BPfold's $H_w = 0.637$ alone exceeds RiNALMo's full RUI = 0.524. This is the operational basis for treating RBI and RUI as distinct axes from micro-F1. Sample sizes match Fig. 3.

**Extended Data Table 1. The Shifu Trifecta on the Shifu-Corpus validation split** ($n$ =25412). Same column conventions as Table 1.

| Model | micro-F1 | RBI | RUI | Coverage | $\text{Tail}_{20}$ | Quadrant |
|---|---|---|---|---|---|---|
| ERNIE-RNA | 0.743 | 0.616 | 0.678 | 0.866 | 0.438 | Broad & Aware |
| PriFold | 0.709 | 0.525 | 0.736 | 0.830 | 0.332 | Broad & Aware |
| RiNALMo | 0.563 | 0.465 | 0.476 | 0.866 | 0.250 | Broad & Blind (margin) |
| MXfold2 | 0.540 | 0.557 | 0.721 | 0.865 | 0.359 | Broad & Aware |
| BPfold | 0.324 | 0.304 | 0.590 | 0.999 | 0.093 | Fragile & Aware |
| UFold | 0.191 | 0.000 | 0.351 | 0.956 | 0.000 | Fragile & Blind |

**Extended Data Table 2. Internal model specifications and training.** Backbone, parameter-efficient adaptation, deployed parameter count (full precision; backbone + head with adapters merged), training corpus, curriculum stages, single-GPU training time, and Shifu-Corpus test metrics for each internal checkpoint. The main-text Table 1 features four of these; one further checkpoint is listed here for completeness: Shifu-LMR-B, a v0-backbone variant with higher-rank adapters. Training times marked * are log-span upper bounds (include idle/resume); LMR-bpRNA is a clean run; the training device was not recorded. The geometric model's Shifu-Corpus metrics are from the DoRA-aware re-evaluation. Values: `model_params.json` (parameters), `exact_model_metrics.csv` (metrics), and the checkpoint configurations/logs (training). All structural models share the recipe: frozen backbone + LoRA + axial pairing head + nested-DP decoder, curriculum stages S0 (Rfam soft-label warm-up), S1 (pairing init), S2 (joint), S3 (joint + auxiliary heads), S4 (hard-negative mining), at max length 512 and effective batch 32 in `bfloat16`.

| Model | Backbone | Adapt. | Params | Corpus | Curric. | GPU-h | micro-F1 | RBI | RUI |
|---|---|---|---|---|---|---|---|---|---|
| Shifu-LMR-Nano | LMR-Nano (64 M) | LoRA r64 | 65 M | Shifu-Corpus | S0–S4 | ∼230* | 0.789 | 0.580 | 0.841 |
| Shifu-LMR-B | LMR v0 (289 M) | LoRA r32 | 291 M | Shifu-Corpus | S0–S4 | ∼217* | 0.775 | 0.573 | 0.852 |
| Shifu-LMR-v0 | LMR v0 (289 M) | LoRA r16 | 291 M | Shifu-Corpus | S0,S2–S4 | ∼231* | 0.776 | 0.591 | 0.855 |
| LMR-bpRNA | LMR v0 (289 M) | LoRA r16 | 291 M | bpRNA-1m | S0,S2–S4 | ∼38 | 0.647 | 0.607 | 0.846 |
| Shifu-LMR-G | LMR-G (86 M) | LoRA r32+DoRA | 88 M | Shifu-Corpus | S0–S4 | ∼269* | 0.787 | 0.521 | 0.810 |

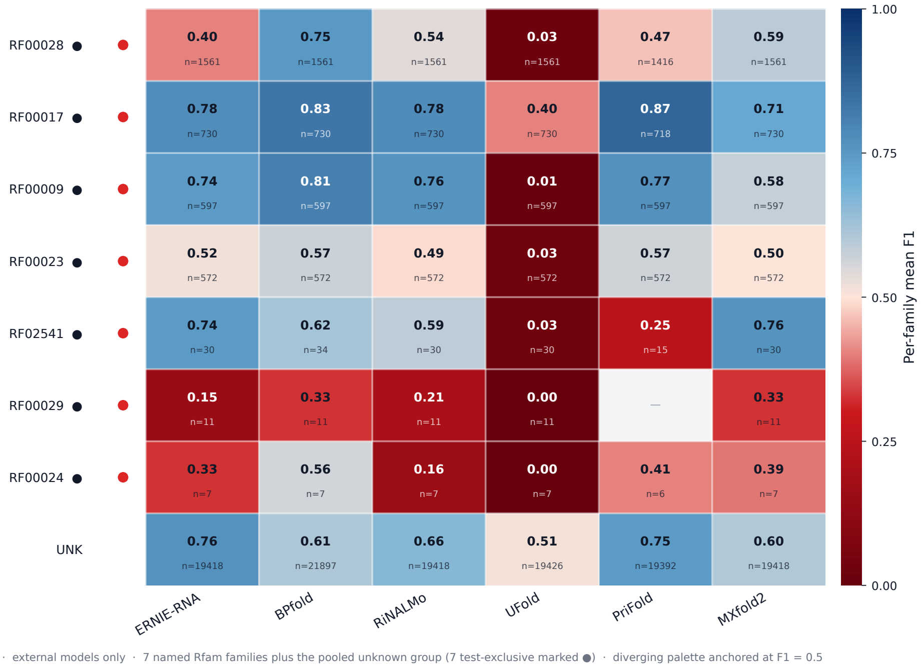


**a.** Shifu-Corpus test (7 named Rfam families plus the pooled *unknown* group; the 7 test-exclusive families marked •).

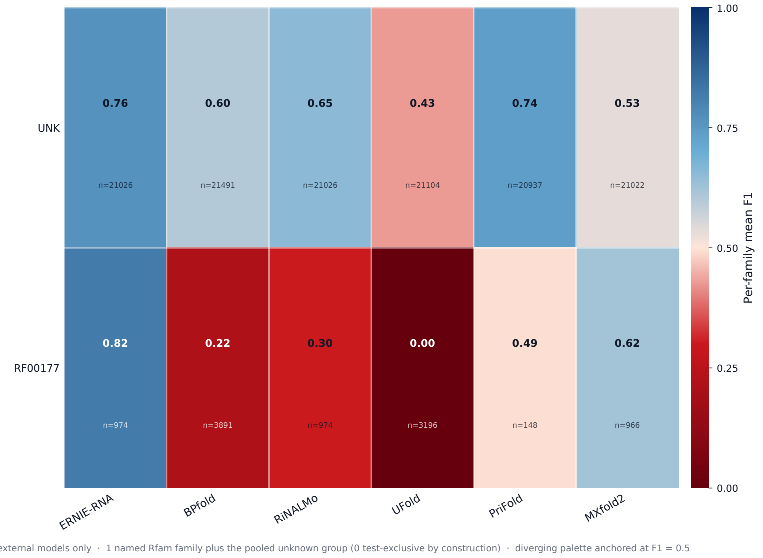


**b.** Shifu-Corpus val (1 named Rfam family plus the pooled *unknown* group; 0 test-exclusive in val by construction).

**Extended Data Fig. 3. Per-Rfam family performance.** Per-family mean micro-F1 for the six external models on Shifu-Corpus test (**a**) and validation (**b**). The seven test-exclusive Rfam families (RF00028, RF00017, RF00009, RF00023, RF02541, RF00029, RF00024; marked • in panel **a**) provide a family-level view of cross-family generalization that complements the (source, length-bin) view in Extended Data Fig. 1. PriFold reports its strongest performance on RF00017, RF00009, and RF00023 (the bpRNA-shaped held-out families); MXfold2 reports its strongest performance on RF02541; BPfold reports its strongest performance on RF00028 and RF00029. No single model performs best on every held-out family, which is consistent with the bpnew result in Fig. 1. Cell labels show per-family mean F1 and unit $n$; the dash on PriFold/RF00029 marks a unit the model could not process. Sample sizes match Fig. 3; per-family $n$ is reported within each cell.

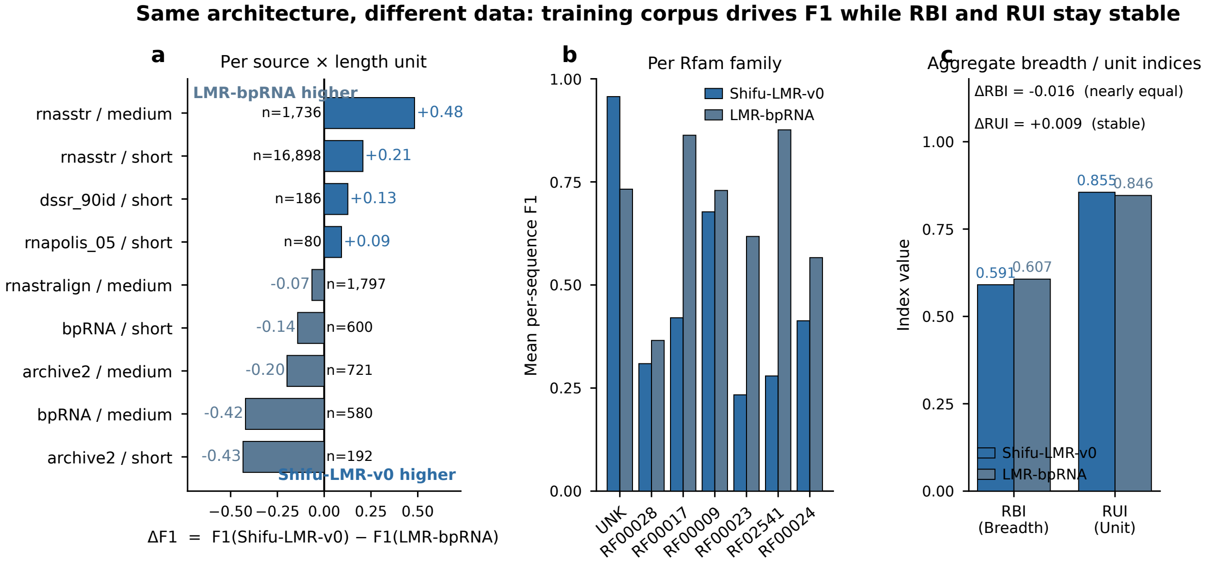


**Extended Data Fig. 4. Same architecture, different training data.** Two models with an identical architecture (LMR v0 backbone + low-rank adapters + axial pairing head + nested dynamic-programming decoder) that differ only in training corpus, on the Shifu-Corpus test split ($n = 25413$). **a**, Per-(source × length) unit micro-F1 difference $\Delta$micro-F1 = micro-F1$_{\text{Shifu-LMR-v0}}$ − micro-F1$_{\text{LMR-bpRNA}}$; blue marks units where the Shifu-trained model wins, orange where the bpRNA-trained model wins, each annotated with unit $n$. **b**, Family-level mean micro-F1 for both models across Rfam families. **c**, Aggregate indices: the training corpus moves micro-F1 (Shifu-LMR-v0 0.776 vs LMR-bpRNA 0.647) while RBI (0.591 vs 0.607) and RUI (0.855 vs 0.846) stay nearly constant. The corpus, not capacity, drives the change. Sample size Shifu-Corpus test $n = 25413$.
