## Supplementary material for "Shifu: an integrated framework for deep learning of RNA secondary structure": SI

### Supplementary Note 1: Shifu-Corpus construction and audit detail

Shifu-Corpus is built from 310034 raw records across six public sources by: (i) exact SHA-256 deduplication (55641 removed); (ii) a 20–8192 nt length-band filter (final 254123 records); (iii) DSSR canonical-pair normalization with multilevel pseudoknot parsing; (iv) MinHash near-duplicate clustering (6-mers, 128 permutations, Jaccard $\geq 0.90$); (v) tiered Rfam/clan labeling via Infernal `cmscan` (`--cut_ga`); and (vi) family-aware 80/10/10 splitting with a $2\times$ source-mix penalty, certified to zero exact and zero cluster leaks across all split pairs. The full audit suite (`global_summary.json`, `split_summary.json`, `dedup_report.json`, `leakage_report.json`, `label_coverage.` `manifest_hashes.json`) is released with the dataset.

### Supplementary Table S1: Per-source and per-split statistics

Supplementary Table S1. Post-deduplication source composition and the realized split sizes of Shifu-Corpus.

| Source (post-dedup) | *n* |
|---|---:|
| rnasstr | 195368 |
| bpRNA | 27699 |
| rnastralign | 27328 |
| archive2 | 2204 |
| dssr_90id | 1006 |
| RNAsolo (rnapolis_05) | 518 |
| Train / Val / Test | 203298 / 25412 / 25413 |

### Supplementary Table S2: Test-split family composition

Supplementary Table S2. Test-split family composition of Shifu-Corpus. Seven Rfam families are held out of training and validation entirely (test-exclusive); all remaining sequences, predominantly unlabeled at the Rfam level, form the pooled *unknown* group. Counts are from the released split manifests.

| Rfam ID | Family | Test sequences |
|---|---|---:|
| RF00028 | Group I catalytic intron | 1561 |
| RF00017 | Metazoan signal recognition particle RNA | 730 |
| RF00009 | Nuclear RNase P | 597 |
| RF00023 | Transfer-messenger RNA (tmRNA) | 572 |
| RF02541 | Bacterial large-subunit (23S) rRNA | 34 |
| RF00029 | Group II catalytic intron | 11 |
| RF00024 | Telomerase RNA | 7 |
| *unknown* | Sequences without a confident Rfam assignment | 21901 |

### Supplementary Table S3: One-nucleotide shift-tolerance sensitivity

Supplementary Table S3. An exploratory robustness check outside the Trifecta's scope. Exact versus one-nucleotide-tolerant (slip-1) micro-F1 on the Shifu-Corpus test split. slip-1 credits a predicted pair whose two partners are each within one nucleotide of a true pair. Tolerance raises every model by 0.01 to 0.06 and leaves the top of the ranking and the paper's conclusions unchanged, though it can reorder closely-spaced models; this confirms that the all-or-nothing property is a real limitation that does not alter the conclusions. Three models (PriFold, UFold, BPfold) are omitted because their released pseudoknot-stripped dot-brackets under-represent their scored pairs.

| Model | exact micro-F1 | slip-1 micro-F1 |
|---|---|---|
| Shifu-LMR-Nano | 0.789 | 0.805 |
| Shifu-LMR-G | 0.787 | 0.800 |
| Shifu-LMR-v0 | 0.776 | 0.795 |
| ERNIE-RNA | 0.648 | 0.690 |
| RiNALMo | 0.627 | 0.647 |
| MXfold2 | 0.559 | 0.581 |
| EternaFold | 0.531 | 0.553 |
| RNAfold | 0.460 | 0.481 |

### Supplementary Note 2: Metric and entropy derivations

Formal definitions of micro-F1, the RNA Breadth Index (RBI), and the RNA Uncertainty Index (RUI) (including the per-residue entropy estimator, the selective-trust curve, and the capacity-discount rule) are given in the main-text Methods. The staged-curriculum hyperparameters and the exact per-model schedules accompany the code release, so the structure-prediction models can be reproduced from the released data and recipe.

### Supplementary Figure S1: Shifu-LMR training schematics

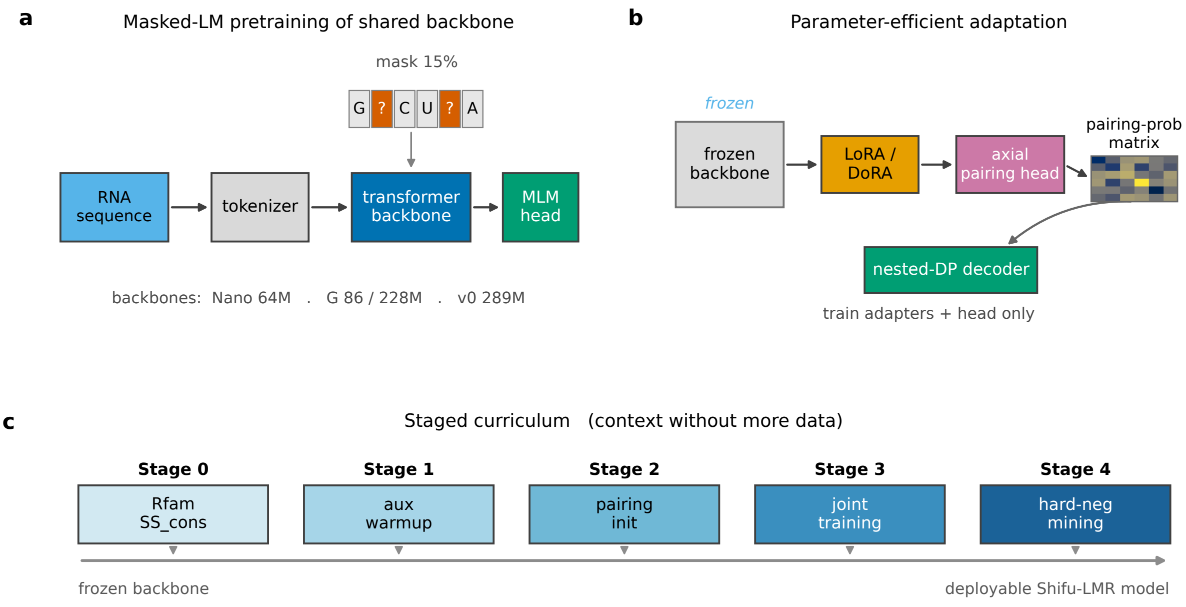


Supplementary Figure S1. Architecture and training of the Shifu-LMR family. **a**, masked-language-model pretraining of a shared RNA backbone (the Nano 64 M, G 86/228 M and v0 289 M backbones). **b**, parameter-efficient adaptation: a frozen backbone with low-rank adapters (LoRA, with DoRA for the geometric backbone), an axial pairing head that emits a symmetric pairing-probability matrix, and a nested dynamic-programming decoder; only the adapters and head are trained. **c**, the staged curriculum (Stage 0 Rfam soft-label warm-up through Stage 4 hard-negative mining) that adds biological context without new data. These schematics accompany the quantitative headline in main-text Fig. 5.
